# bmotif: a package for motif analyses of bipartite networks

**DOI:** 10.1101/302356

**Authors:** Benno I. Simmons, Michelle J. M. Sweering, Maybritt Schillinger, Lynn V. Dicks, William J. Sutherland, Riccardo Di Clemente

## Abstract

1. Bipartite networks are widely-used to represent a diverse range of species interactions, such as pollination, herbivory, parasitism and seed dispersal. The structure of these networks is usually characterised by calculating one or more metrics that capture different aspects of network architecture. While these metrics capture useful properties of networks, they only consider structure at the scale of the whole network (the macro-scale) or individual species (the micro-scale). ‘Meso-scale’ structure between these scales is usually ignored, despite representing ecologically-important interactions. Network motifs are a framework for capturing this meso-scale structure and are gaining in popularity. However, there is no software available in R, the most popular programming language among ecologists, for conducting motif analyses in bipartite networks. Similarly, no mathematical formalisation of bipartite motifs has been developed.
2. Here we introduce bmotif: a package for counting motifs, and species positions within motifs, in bipartite networks. Our code is primarily an R package, but we also provide MATLAB and Python code of the core functionality. The software is based on a mathematical framework where, for the first time, we derive formal expressions for motif frequencies and the frequencies with which species occur in different positions within motifs. This framework means that analyses with bmotif are fast, making motif methods compatible with the permutational approaches often used in network studies, such as null model analyses.
3. We describe the package and demonstrate how it can be used to conduct ecological analyses, using two examples of plant-pollinator networks. We first use motifs to examine the assembly and disassembly of an Arctic plant-pollinator community, and then use them to compare the roles of native and introduced plant species in an unrestored site in Mauritius.
4. bmotif will enable motif analyses of a wide range of bipartite ecological networks, allowing future research to characterise these complex networks without discarding important meso-scale structural detail.

## Introduction

Bipartite networks are widely used to study the structure of interactions between two groups of species, such as plants and pollinators, hosts and parasitoids, and plants and seed dispersers (Borrett, Moody, & Edelmann, 2014). Studies of bipartite networks have yielded many new insights (Bascompte & Jordano, 2007), such as uncovering widespread nestedness and modularity in mutualistic communities (Bascompte, Jordano, Melián, & Olesen, 2003; Olesen, Bascompte, Dupont, & Jordano, 2007), and showing that community structure is stable despite substantial turnover in species and interactions over space and time (Petanidou, Kallimanis, Tzanopoulos, Sgardelis, & Pantis, 2008; Dáttilo, Guimarães, & Izzo, 2013). Such studies typically describe networks with one or more metrics, such as connectance (the proportion of possible interactions which are realised), nestedness (the extent to which specialist species interact with subsets of the species generalist species interact with), degree (number of partners a species has) and *d′* (the extent to which a species’ interactions deviate from a random sampling of its partners).

While these metrics describe useful properties of networks, macro-scale measures, such as connectance and nestedness, can be too broad to capture fine-scale details, while micro-scale metrics, such as degree as *d′*, can be too narrow to capture a species’ indirect interactions (Cirtwill, Roslin, Rasmussen, Olesen, & Stouffer, 2018). Capturing network structure at the meso-scale between these macro and micro scales is necessary to overcome these issues (Cirtwill et al., 2018). For example, a micro-scale metric such as degree might show that a plant is visited by two pollinators, while meso-scale structure could reveal that one of these pollinators is a generalist visiting three other generalist plants, while the other is a specialist visiting only the focal plant. Such distinctions can have important consequences for understanding the ecology and evolution of communities and so are essential to incorporate in network analyses.

To capture meso-scale structure, ecologists are increasingly using bipartite motifs: subnetworks representing interactions between a given number of species (Fig. 1). These subnetworks can be thought of as the basic ‘building blocks’ of networks (Milo et al., 2002). Bipartite motifs are used in two main ways. First, to calculate how frequently different motifs occur in a network. For example, Rodríguez-Rodríguez et al. (2017) used this approach to show that plant species involved in both mutualistic and antagonistic interactions with animals were the most important for pollination. Second, to quantify species roles in a community by counting the frequency with which species occur in different positions within motifs. For example, Baker et al. (2015) used this method to demonstrate that species’ roles in host-parasitoid networks are an intrinsic property of species. However, while the motif framework is gaining in popularity, no software currently exists to conduct motif analyses of bipartite networks in R, the most popular programming language among ecologists.

**Figure 1:**
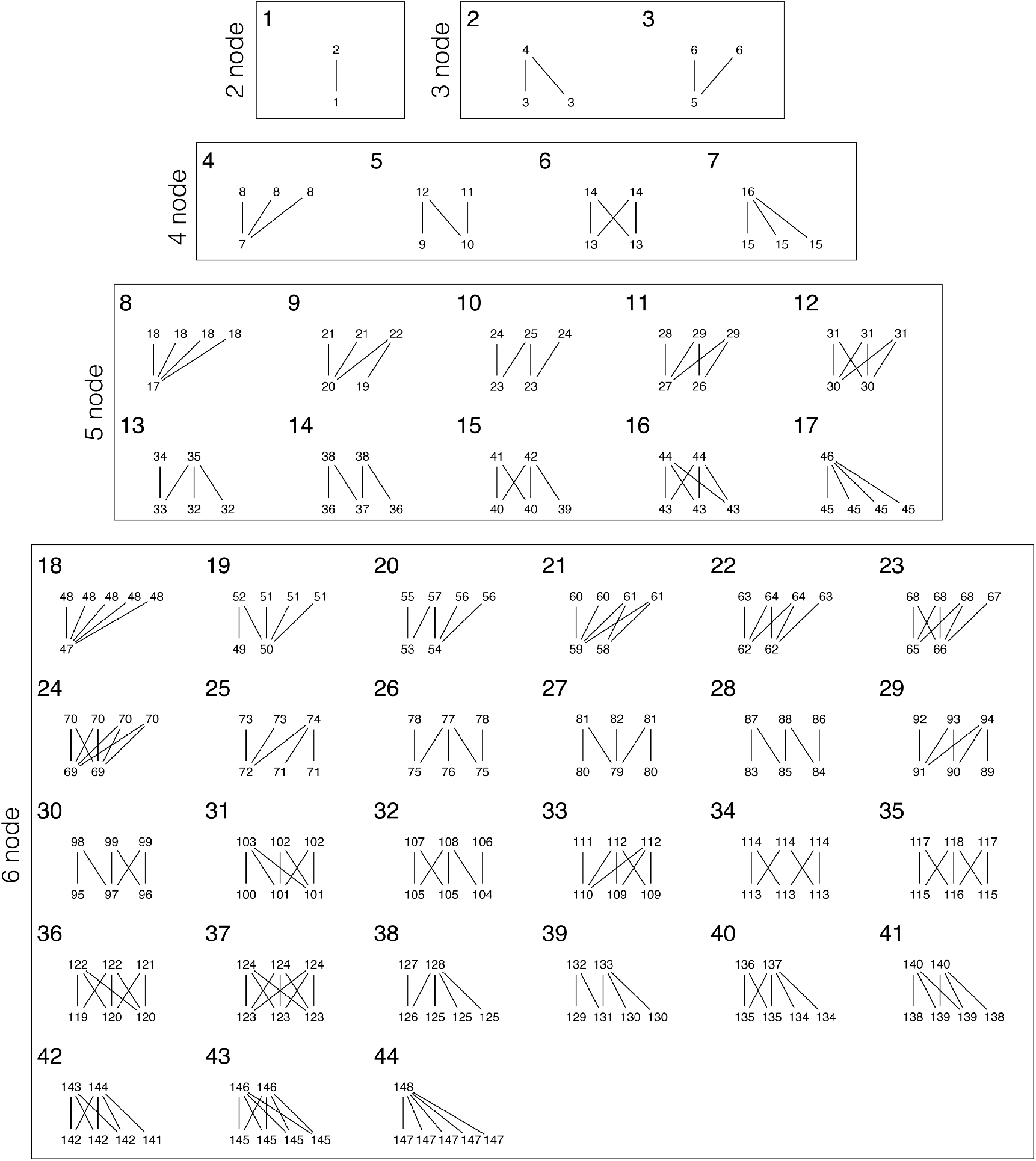
All bipartite motifs containing up to 6 nodes (species). Large numbers identify each motif. Small numbers represent the unique positions species can occupy within motifs, following Baker et al. (2015) Appendix 1. Lines between small numbers indicate undirected species interactions. There are 44 motifs containing 148 unique positions.

To fill this gap, we introduce bmotif: an R package, based on a formal mathematical framework, for counting motifs, and species positions within motifs, in bipartite networks. While bmotif is primarily an R package, we additionally provide MATLAB and Python code that replicates the core package functionality. Here, we introduce the motifs and motif positions counted by bmotif and describe the package’s main functions and performance. We then provide two examples showing how bmotif can be used to answer questions about ecological communities. While here we focus on mutualistic bipartite networks, our methods are general and can also be applied to other types of interaction, such as parasitism and herbivory, and even non-biological systems, such as trade networks (Saracco, Di Clemente, Gabrielli, & Squartini, 2015) and finance networks (Gualdi, Cimini, Primicerio, Di Clemente, & Challet, 2016).

## Description

### Defining bipartite motifs

In a bipartite network containing *N* species, a motif is a subnetwork comprising *n* species and their interactions (where *n* < *N* and all species have at least one interaction). Fig. 1 shows the motifs included in bmotif: all 44 possible motifs containing up to six nodes. Within motifs, species can appear in different positions (Fig. 1). For reasons of symmetry, not all these positions are topologically unique (Stouffer, Sales-Pardo, Sirer, & Bascompte, 2012). For example, motif six contains four species, but only two positions (Fig. 1). The 148 unique positions a species can occupy across all motifs up to six nodes are shown in Fig. 1. Motifs and positions are ordered as in Baker et al. (2015) Appendix 1.

Networks in bmotif are represented as incidence matrices (**M**), with one row for each species in the first set (such as pollinators) and one column for each species in the second set (such as plants). When species *i* and *j* interact, m*_ij_* = 1; if they do not interact m*_ij_* = 0. This widely-used representation was chosen for compatibility with other packages (Dormann, Frund, Bluthgen, & Gruber, 2009) and open-ccess network repositories, such as the Web of Life (www.web-of-life.es). Species in rows correspond to nodes in the top level of the motifs in Fig. 1; species in columns correspond to nodes in the bottom level.

### Main functions

bmotif has two functions: (i) *mcount,* for calculating how frequently different motifs occur in a network, and (ii) *positions,* for calculating the frequency with which species (nodes) occur in different positions within motifs to quantify a species’ structural role. To enumerate motif frequencies and species position counts, bmotif uses mathematical operations directly on the incidence matrix: for the first time, we derive 44 expressions for each of the 44 motifs and 148 expressions for each of the 148 positions within motifs (Appendix S2).

*mcount* takes a network as input and returns a data frame with one row for each motif (17 or 44 rows depending on whether motifs up to five or six nodes are requested, respectively) and three columns. The first column is the motif identity as in Fig. 1; the second column is the motif size class (number of nodes each motif contains); and the third column is the frequency with which each motif occurs in the network (a network’s motif profile). For comparing multiple networks it is important to normalise motif frequencies. Therefore, if the ‘normalise’ argument is TRUE, three columns are added to the data frame, each corresponding to a different method for normalising motif frequencies. The first column (‘normalise_sum’) expresses the frequency of each motif as a proportion of the total number of motifs in the network. The second column (‘normalise_sizeclass’) expresses the frequency of each motif as a proportion of the total number of motifs within its size class. The final column (‘normalise_nodesets’) expresses the frequency of each motif as the number of species combinations that occur in a motif as a proportion of the number of species combinations that could occur in that motif. For example, in motifs 9, 10, 11 and 12, there are three species in the top set (*A*) and two species in the lower set (*B*) (Fig. 1). Therefore, the maximum number of species combinations that could occur in these motifs is given by the product of binomial coefficients, choosing three species from *A* and two from 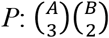 (Poisot & Stouffer, 2016). The most appropriate normalisation depends on the question being asked. For example, ‘normalise_sum’ allows for consideration of whether species are more involved in smaller or larger motifs. Conversely, ‘normalise_sizeclass’ focuses the analysis on how species form their interactions among different arrangements of *n* nodes.

*positions* takes a network as input and returns a data frame, **W**, with one row for each species and one column for each position (46 or 148 columns, depending on whether motifs up to five or six nodes are requested, respectively; Fig. 1). *W_rc_* gives the number of times species *r* occurs in position c. Each row thus represents the structural role or ‘interaction niche’ of a species. The ‘level’ argument allows positions to be requested for all species, species in set *A* only or species in set *B* only, returning a data frame with *A* + *B* rows, *A* rows or *B* rows, respectively. Two types of normalisation are provided: ‘sum’ normalisation expresses a species’ position frequencies as a proportion of the total number of times that species appears in any position; ‘size class’ normalisation uses the same approach, but normalises frequencies within each motif size class. Again, the most appropriate normalisation depends on the question being asked: if movements between motif size classes are of interest, ‘sum’ normalisation is most appropriate; if the focus is on how species form interactions among a given number of nodes, then ‘size class’ normalisation should be chosen.

## Computational performance

To assess the speed of bmotif functions, we used *mcount* and *positions* to calculate the complete motif profiles of 175 empirical pollination and seed dispersal networks and the positions of all their constituent species. While most of these networks use the frequency of animal visits to plants as a surrogate for true pollination or seed dispersal, this has been shown to be a reasonable proxy (Vázquez, Morris, & Jordano, 2005; Simmons et al., 2018). The networks varied in size from 6 to 797 species (mean: 77.1; standard deviation: 117.8). Further details of the networks used for this analysis are given in Supplementary Data 1. Analyses were carried out on a computer with a 4.0 GHz processor and 32 GB of memory. Functions were timed using the R package ‘microbenchmark’ (Mersmann, 2015). Results are shown in Fig. 2.

**Figure 2:**
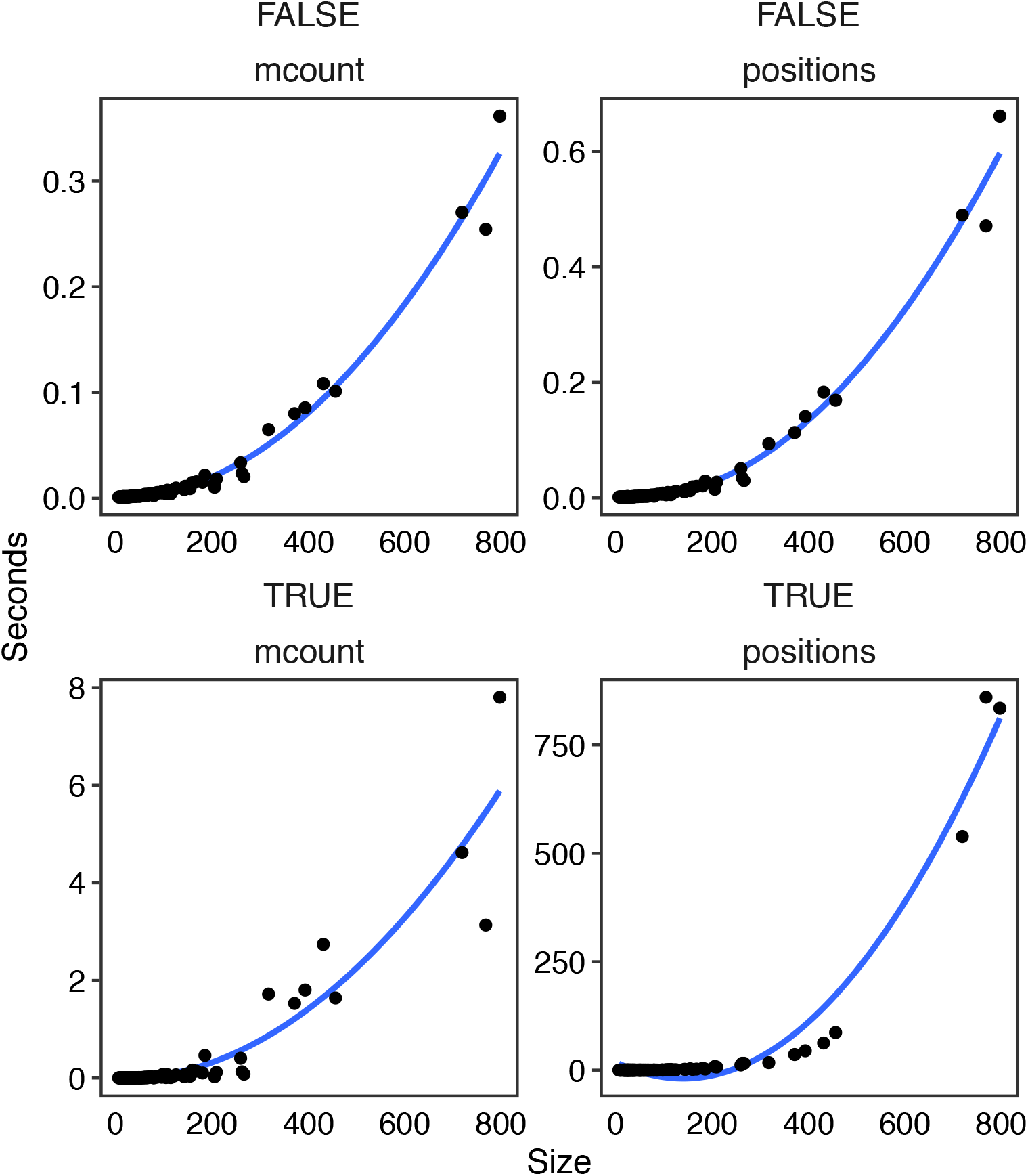
Relationship between network size and computational performance for *mcount* and *positions.* Functions were timed on 175 empirical networks, for motifs containing up to five and six nodes. Lines are best fit polynomial curves of degree 2.

As expected, the time taken for a function to run increases monotonically with the size of the network (number of species). When six-node motifs were excluded, *mcount* and *positions* took 0.36 and 0.66 seconds, respectively, to complete for the largest network in our dataset (797 species). For smaller networks which are more typical of the communities analysed by ecologists, both functions completed in substantially less than one second. This speed is possible as all formulae involved in calculations of motifs up to five-nodes use relatively simple operations, such as matrix multiplication or the binomial coefficient. When six-node motifs were included, for a network with 78 species (close to the mean network size of 77.1 species), *mcount* completed in 0.01 seconds, while *positions* completed in 0.32 seconds. For the largest network, *mcount* completed in 7.8 seconds, while *positions* took 13.9 minutes. Six-node motifs slow down calculations as, unlike five-node motifs, their algorithms require the use of the tensor product.

We carried out additional analyses using randomly-generated networks to disentangle the effects of both network dimensions and connectance on computational performance (Appendix S1). We found that connectance had little effect on the performance of individual motif and position calculations, while a polynomial of degree two explained the increase in time with network size (R^2^ > 0.99) (Appendix S1).

## Example analyses

### Comparing community structures

Here we use bmotif to examine the assembly and disassembly of an Arctic plant-pollinator community. Networks were sampled daily, when weather conditions allowed, at the Zackenberg Research Station in northeastern Greenland, across two full seasons in 1996 (24 days) and 1997 (26 days) (Olesen, Bascompte, Elberling, & Jordano, 2008). Basic network properties are given in Supplementary Data 2. We used *mcount* to calculate motif frequencies in each daily network in both years, normalised using ‘normalise_nodesets’. Days 1 and 24 in 1996, and days 1 and 26 in 1997, were excluded from the analysis as they were too small for some motifs to occur. Using nonmetric multidimensional scaling (NMDS), we visualised how the community structure changed from assembly after the last snow melt to disassembly at the first snow fall, in two consecutive years (Fig. 3). More positive values of the first NMDS axis are associated with motifs where generalist pollinators compete for generalist plants, while negative values are associated with motifs where more specialist pollinators have greater complementarity in the specialist plants they visit. More positive values of the second NMDS axis are associated with loosely connected motifs containing specialist plants interacting with both specialist and generalist pollinators, while negative values are associated with highly connected motifs containing pollinators competing for generalist plants. While the community was relatively stable over time in the 1996 season, there were larger structural changes in 1997, with a largely monotonic shift from high competition between generalist pollinators at the start of the season, to lower competition between more specialist pollinators at the end of the season, with a more complementary division of plant resources (Fig. 3). Thus while network structure may appear stable when analysed with traditional indices such as connectance (Olesen et al., 2008), motifs reveal the presence of complex, ecologically-important structural dynamics. Additionally, it is clear that, even in consecutive years, the community followed different structural trajectories, emphasising the danger of treating networks as static entities (Rasmussen, Dupont, Mosbacher, Trøjelsgaard, & Olesen, 2013).

**Figure 3:**
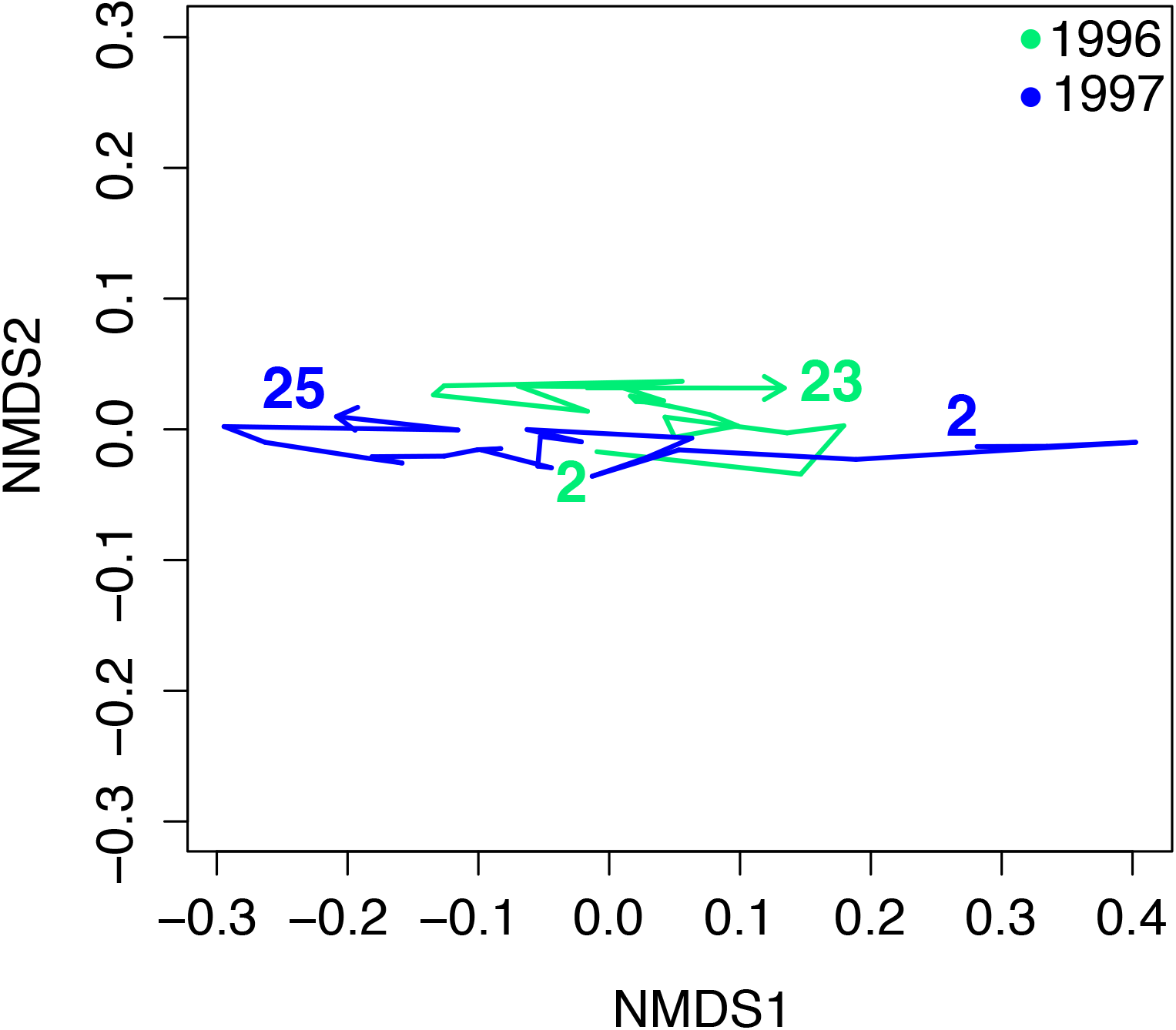
Nonmetric multidimensional scaling plot (NMDS) showing change in Arctic plant-pollinator network structure over the 1996 and 1997 seasons, quantified using motifs. Numbers represent the days of sampling.

### Comparing species’ structural roles

We use *positions* to compare the roles of native and introduced plant species in a plant-pollinator community sampled in Mauritius in November 2003 (Kaiser-Bunbury, Memmott & Müller 2009; 48 species, 75 interactions, connectance = 0.134). We calculated the sum-normalised roles of all plant species (16 native and 4 introduced) and plotted them on two NMDS axes (Fig. 4). This figure shows three striking features. First, there is almost no overlap between native and introduced species’ interaction niches. Similar to research showing that non-native species can occupy different functional niches to native species (Ordonez, Wright, & Olff, 2010), these results suggest they may also occupy unexploited interaction niches. Further research could use motifs to investigate whether introduced species ‘pushed’ native species out of previously occupied interaction niche space, or whether introduced species colonised previously-unused space. Second, the interaction niche of introduced species is much smaller than that of native species: the four introduced species all occupy similar areas of motif space, possibly suggesting a single ‘invader role’. Third, introduced species occupy lower values on the second NMDS axis, corresponding to motif positions where they are visited by generalist pollinator species, possibly due to the absence of co-evolutionary associations with specialists. NMDS analyses were conducted with the metaMDS function in the R package vegan (Oksanen et al., 2016).

**Figure 4:**
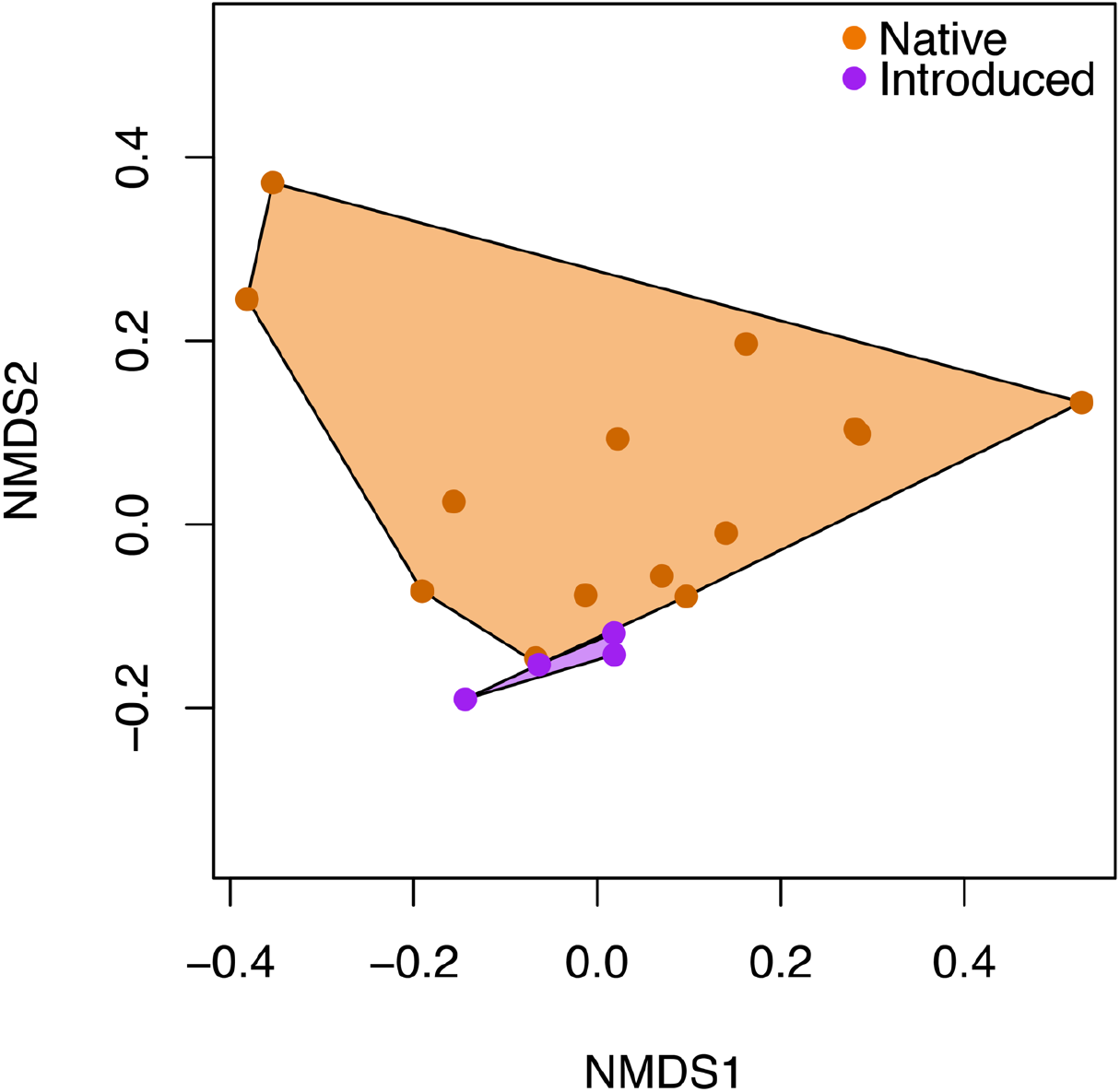
The roles of native and introduced species in a plant-pollinator network. Each point represents the role of a species in the network. Shaded polygons are convex hulls either containing all introduced species or all alien species.

## Implementation and availability

The bmotif package is available for the R programming language. The package can be installed in R using install.packages(“bmotif’). This paper describes version 1.0.0 of the software. The source code of the package is available at https://github.com/SimmonsBI/bmotif-release. Any problems can be reported using the *Issues* system. The code is version controlled with continuous integration and has code coverage of approximately 98%. MATLAB and Python code replicating the core package functionality is available at https://github.com/SimmonsBI/bmotif-matlab and https://github.com/SimmonsBI/bmotif-python respectively. All code is released under the MIT license.

## Conclusions

bmotif is an R package and set of mathematical formulae enabling motif analyses of bipartite networks. Specifically, bmotif provides functions for two key analyses: (i) enumerating the frequency of different motifs in a network, and (ii) calculating how often species occur in each position within motifs. These two techniques capture important meso-scale variation in network structure that may be missed by traditional methods. Motif approaches represent a new addition to the network ecologists ‘toolbox’ for use alongside other techniques to analyse bipartite networks. We hope bmotif encourages further uptake of the motif approach to shed light on the ecology and evolution of species and communities.

## Acknowledgements

BIS is supported by the Natural Environment Research Council as part of the Cambridge Earth System Science NERC DTP [NE/L002507/1]. BIS, MJM and MS acknowledge the Cambridge Faculty of Mathematics Bridgwater Summer Research Fund/CMP bursary fund for support. LVD is funded by the Natural Environment Research Council (grant code: NE/N014472/1). WJS is funded by Arcadia. RDC as Newton International Fellow of the Royal Society acknowledges support from The Royal Society, The British Academy and the Academy of Medical Sciences (Newton International Fellowship, NF170505).

## Author contributions

BIS conceived the project, conducted analyses and wrote the first draft of the manuscript. BIS, MJM and RDC developed the bmotif package. BIS, WJS, LVD and RDC planned the study. BIS and RDC coordinated and designed the work. All authors contributed to writing the manuscript.

## Data accessibility

All networks used in this study are available from the Web of Life repository (www.web-of-life.es), with the exception of the Greenland plant-pollinator networks which are available from Data Dryad (Saavedra et al., 2016)

